# Assessing Fairness of AlphaFold2 Prediction of Protein 3D Structures

**DOI:** 10.1101/2023.05.23.542006

**Authors:** Usman Abbas, Jin Chen, Qing Shao

**Affiliations:** Chemical & Materials Engineering, University of Kentucky, Lexington, Kentucky, USA; Institute for Biomedical, Informatics, University of Kentucky, Lexington, Kentucky, USA

**Keywords:** AlphaFold, protein structures, AI fairness

## Abstract

AlphaFold2 is reshaping biomedical research by enabling the prediction of a protein’s 3D structure solely based on its amino acid sequence. This breakthrough reduces reliance on labor-intensive experimental methods traditionally used to obtain protein structures, thereby accelerating the pace of scientific discovery. Despite the bright future, it remains unclear whether AlphaFold2 can uniformly predict the wide spectrum of proteins equally well. Systematic investigation into the fairness and unbiased nature of its predictions is still an area yet to be thoroughly explored. In this paper, we conducted an in-depth analysis of AlphaFold2’s fairness using data comprised of five million reported protein structures from its open-access repository. Specifically, we assessed the variability in the distribution of PLDDT scores, considering factors such as amino acid type, secondary structure, and sequence length. Our findings reveal a systematic discrepancy in AlphaFold2’s predictive reliability, varying across different types of amino acids and secondary structures. Furthermore, we observed that the size of the protein exerts a notable impact on the credibility of the 3D structural prediction. AlphaFold2 demonstrates enhanced prediction power for proteins of medium size compared to those that are either smaller or larger. These systematic biases could potentially stem from inherent biases present in its training data and model architecture. These factors need to be taken into account when expanding the applicability of AlphaFold2.

## 1 Introduction

Substantial progress has been made in solving the protein folding problem since Anfinsen first posed the question of whether a protein’s three-dimensional (3D) structure can be generated from its sequence alone [1-9]. Gaining a reliable 3D structure is viewed as a fundamental step towards understanding protein functions [2, 4, 7, 8, 10-18], and is especially important in applications like drug design [2, 10, 11]. Traditionally there have been two main approaches, with the first relying mainly on molecular simulations, and the second relying on evolutionary history via bioinformatics methods [10, 19]. However, both paths had failed to compete with the accuracies of experiments until recently with the explosion of artificial intelligence-based models [3, 5-7, 11, 13, 14, 18].

The Critical Assessment of Techniques for Protein Structure Prediction (CASP) competition is widely accepted as the standard for assessing the performance of protein structure prediction models [10, 12, 20, 21]. Given its impressive performance in CASP14 [2, 4, 14, 22-26], AlphaFold2 was the first model to be capable of achieving accuracies comparable to those attained via experiments [9, 15, 19, 24, 25, 27, 28]. CASP uses the Local Distance Difference Test (LDDT) [20] as a numerical measure to evaluate the model performance as it measures the distance between the predicted structure and the ground truth of a protein. This could be done by comparing only the heavy atoms (LDDT) or only the backbone atoms (LDDT-Cα) [10]. AlphaFold2 uses a variant of LDDT-Cα called Predicted Local Distance Difference Test (PLDDT), as a measure of per-residue prediction credibility [10, 14, 29].

AlphaFold2 has displayed remarkable accuracy in predicting 3D structures of proteins [3]. Despite this, it is imperative that blind trust is not ascribed to AlphaFold2 since most models tend to have a scope within which they are most useful [3, 15]. AI model fairness depends on model architecture and training data [3, 30, 31]. However, many users of AlphaFold2 are not experts in deep learning, and this raises concerns regarding when to rely on AlphaFold2’s protein structure predictions. These concerns become more critical as AlphaFold2 is increasingly being used in structure-based drug design and protein functional research where misinterpretations could prove costly [3, 17, 29]. Using Google DeepMind’s criteria [29] for judging PLDDT scores, it is the aim of this work to allay some of those concerns and use the publicly available protein structure predictions from AlphaFold2’s protein structure database [29] of over 200 million proteins to carry out a systematic statistical analysis of the PLDDT scores generated per amino acid.

In this work, we establish a framework to systematically assess the fairness of AlphaFold2, i.e., whether its performance is significantly independent of protein types. The framework includes three steps, i.e., the selection, assessment, and aggregation of predicted 3D structures in large scale. We first classify proteins into multiple groups based on the standard amino acids, secondary structures, and protein sizes, respectively. We then apply statistical analysis on the PLDDT scores of AlphaFold2 predictions in these groups. As per DeepMind’s guidance, the prediction credibility of AlphaFold2 is quantified using the PLDDT score of each individual amino acid. Using PLDDT as a proxy for accuracy, we can observe the variation of prediction confidence, if any, with respect to amino acid types, secondary structure types, and protein size. Finally, we aggregate and compare results from all protein groups. To the best of our knowledge, this kind of large-scale statistical study has not yet been conducted in the literature. The rest of this paper is structured as follows: Section 2 describes our methodology, section 3 presents the results and discussion, and section 4 draws the conclusion.

## 2 Methodology

The five batches, each containing one-million proteins, were randomly selected from the AlphaFold2 repository v4 of over 200 million proteins. Stepwise, we downloaded the full list of all protein IDs from the v4 repository, randomly selected one-million proteins from this list and then downloaded the related PDB files from the repository. The selected proteins are then removed from the list to avoid duplication in subsequent batches. We repeated this process to create five one-million protein batches. The corresponding amino acid sequences and PLDDT were curated from the downloaded PDB files directly. The secondary structure of individual amino acids was calculated based on the hydrogen bond structure using the Define Secondary Structure of Proteins (DSSP) algorithm [32, 33]. The sequence length of individual proteins was calculated based on the total number of amino acids. All these raw data were stored as text files for further statistical analysis using Python-3.10, Pandas, Matplotlib and NumPy packages.

## 3 Results and Discussion

In this section, we present results of statistical analysis of five million protein structure predictions obtained from the AlphaFold2 database. Deepmind uses a set of criteria [29] to categorize AlphaFold2’s predictions into four confidence levels based on the PLDDT scores: (a) high: PLDDT ???????? 90, (b) medium: 70 ≤ PLDDT < 90, (c) low: 50 ≤ PLDDT < 70, and (d) very low: PLDDT < 50. We will use the same four categories in our results and discussion.

### 3.1 The population distribution of amino acids

With the AlphaFold2 v4 database currently containing over 200 million proteins, it is imperative that our sampling across multiple batches is consistent. In Figure 1, we present the population distribution of all 20 amino acid types from the proteins in batch 1. Figure S1 shows the distributions for the other batches. The population of each amino acid type does not change much across the five batches. ALA hovers around 30,000,000, VAL is around 22,000,000 while TRP is around 4,000,000 in population size.

**Figure 1.**
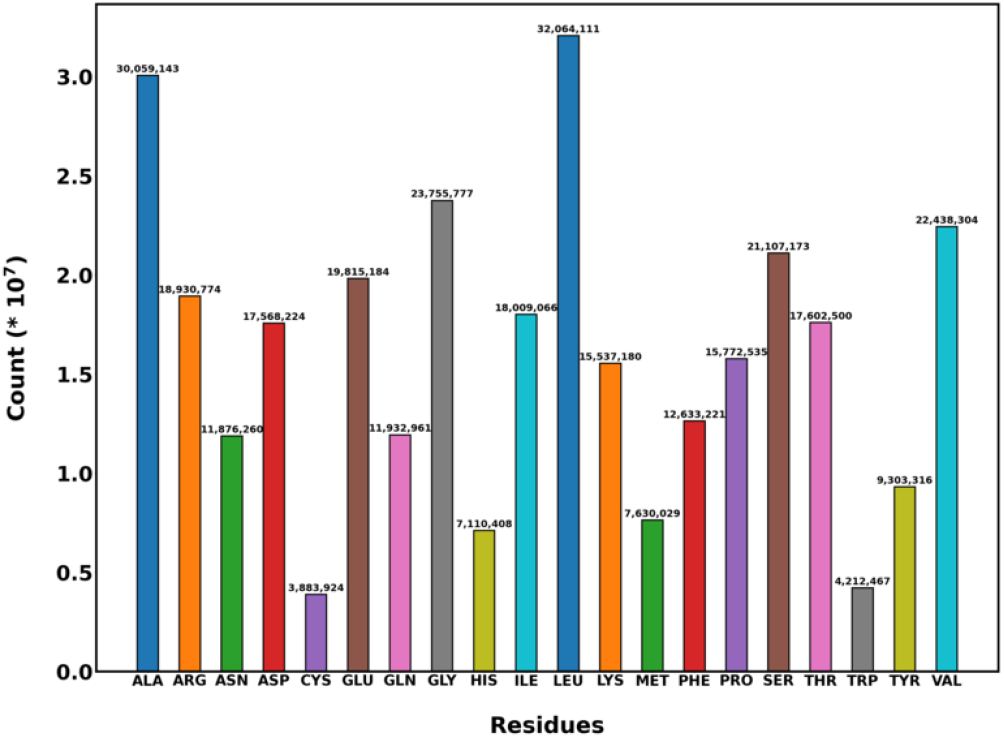
The population distribution of the 20 amino acid types in batch 1.

This consistency is also observed for the population distribution of secondary structures in Figure 2. Figure S2 shows the data for the other batches. The alpha-helix is around 123,000,000 while the coil is around 79,000,000 in population size across all 5 batches.

**Figure 2.**
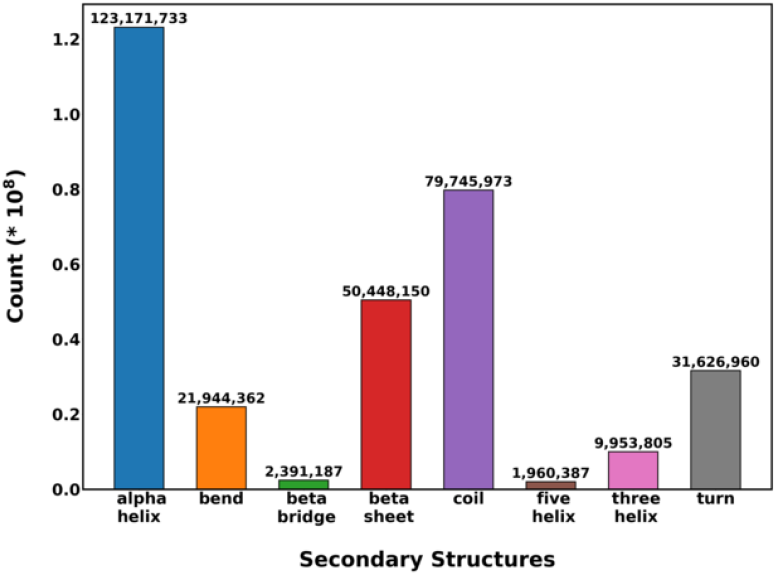
The population distribution of the 8 secondary structures in batch 1.

Figure 3 presents the variation of the population size of amino acid types by protein size N in batch 1. Plots for batches 2-5 can be found in Figures S3 – S7. The relative trends remain the same as that of the overall batches in Figure 1 and S1, but the highest population sizes for each amino acid type are observed outside the extremes of N < 100 and N > 1000 amino acids. For instance, ALA has a population size of about 3,000,000 for proteins with 500 ≤ N < 600 amino acids compared to about 1,000,000 for N > 1000 amino acids.

**Figure 3.**
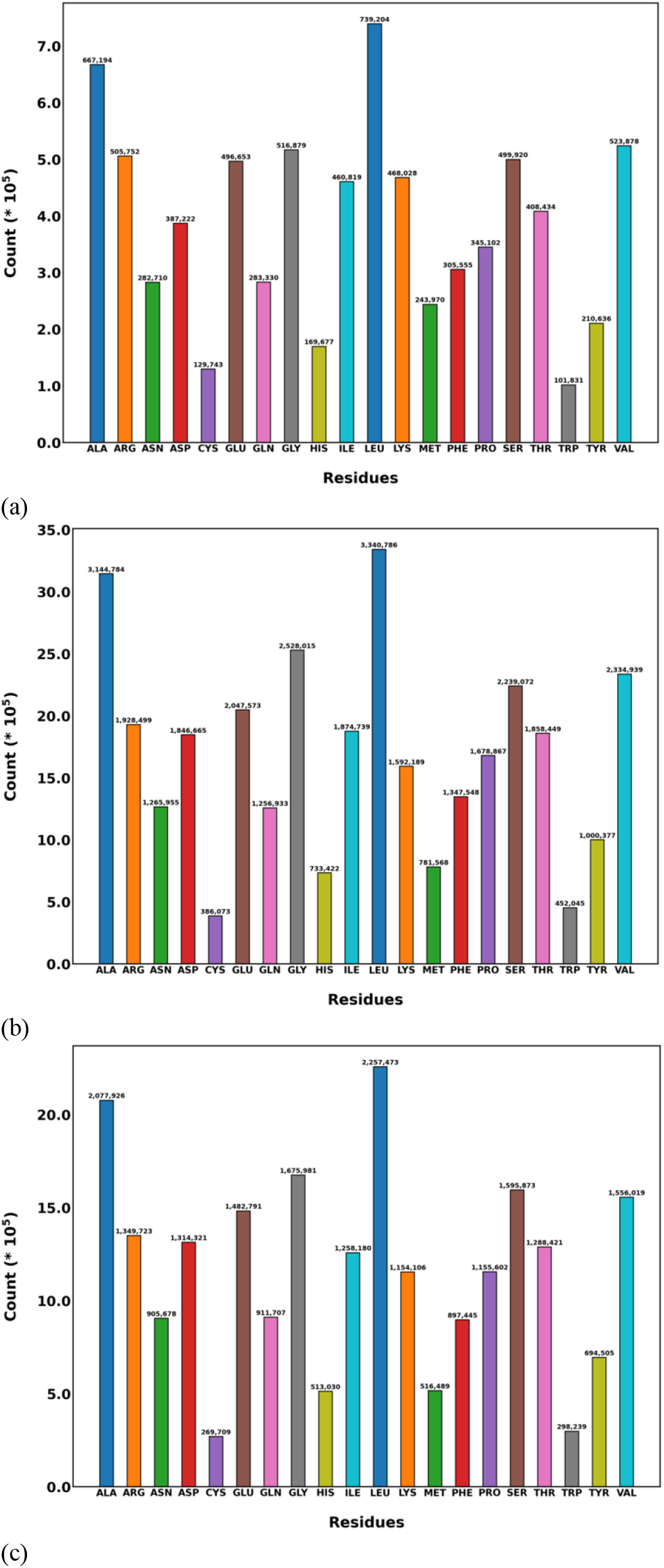
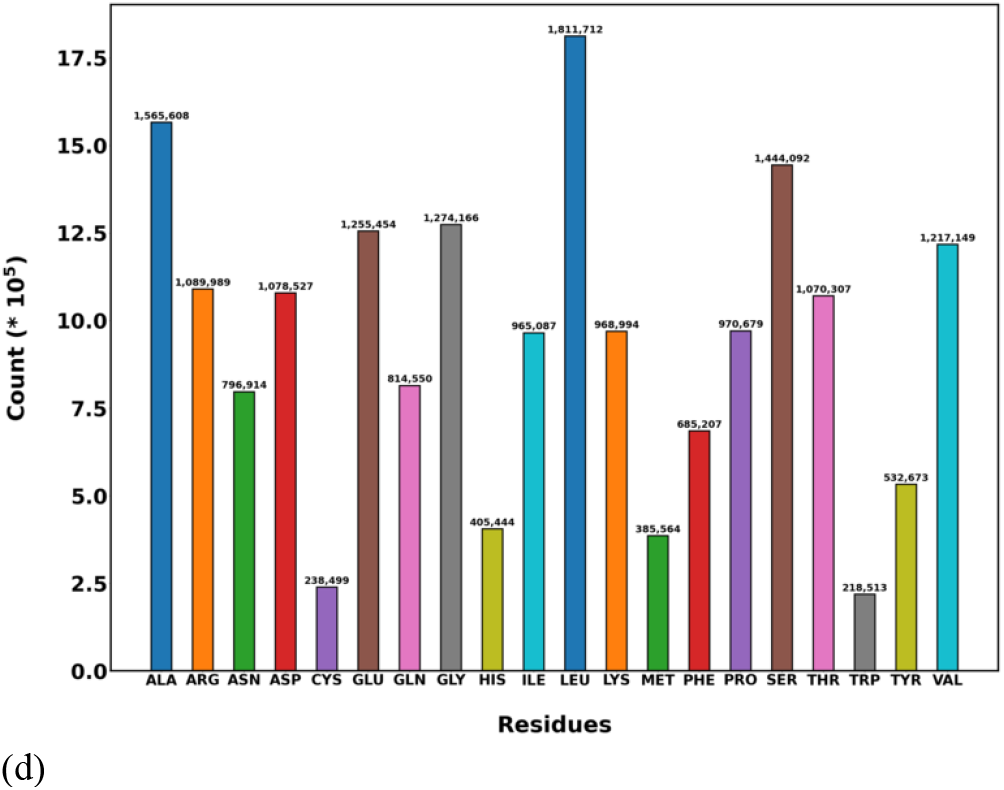
The population distribution of the 20 amino acid types as a function of protein size N in batch 1. (a) N < 100 amino acids, (b) 500 ≤ N < 600 amino acids, (c) 600 ≤ N < 700 amino acids, (d) N > 1000 amino acids.

Figure 4 shows the population distribution of the secondary structure types classified by protein size N in batch 1. Figure S8 – S12 shows the data for the other batches. Across all length classifications, the relative trends for secondary structure population distributions are consistent with the overall batch distributions in Figure 2. Similar to the amino acids in Figure 3, the lowest population sizes for secondary structures are observed in proteins with N < 100 amino acids. The highest population sizes are observed in proteins with N < 100 and N > 1000 amino acids. For instance, the alpha-helix has a population range of 13,000,000 for proteins with 500 ≤ N < 600 amino acids. In contrast, the alpha-helix population drops to about 3,200,000 and 6,000,000 for proteins with N < 100 amino acids and N ≥ 1000 amino acids respectively.

**Figure 4.**
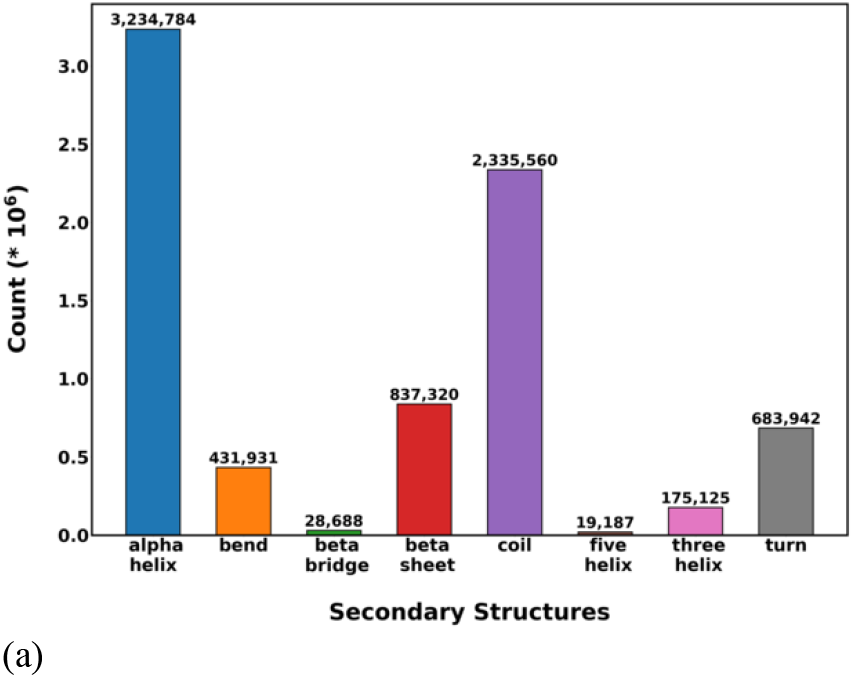
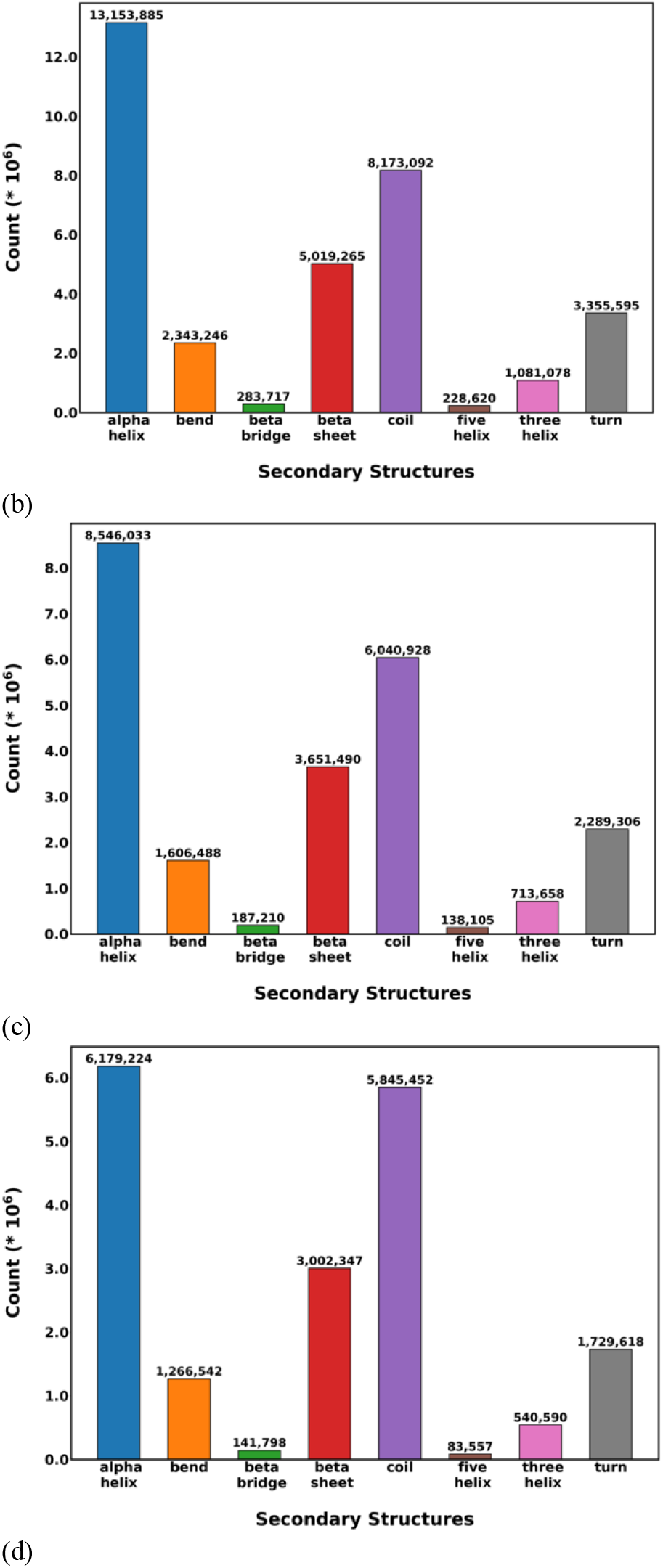
The population distribution of secondary structure types as a function of protein size N in batch 1. (a) N < 100 amino acids, (b) 500 ≤ N < 600 amino acids, (c) 600 ≤ N < 700 amino acids, (d) N > 1000 amino acids.

### 3.2 The variation of prediction credibility with amino acid types

In this section, we analyze the variation of AlphaFold2’s prediction credibility with amino acid types based on the distribution of PLDDT scores. We present the effect of amino acid type on AlphaFold2’s prediction confidence for batch 1 in Figure 5. Plots for other batches can be found in Figure S13. Using DeepMind’s accuracy threshold of PLDDT ≥ 70 for medium model confidence, we assess AlphaFold2’s prediction confidence based on the median PLDDT score for each amino acid. One thing that stands out is the consistency of AlphaFold2’s prediction confidence across all batches. The amino acids with the highest median PLDDT scores in batch 1 are: TRP (94.00), VAL (93.94), ILE (93.88), TYR (93.62), PHE (93.31) and LEU (93.19). The lowest median PLDDT scores are observed for PRO (89.00) and SER (88.38).

**Figure 5.**
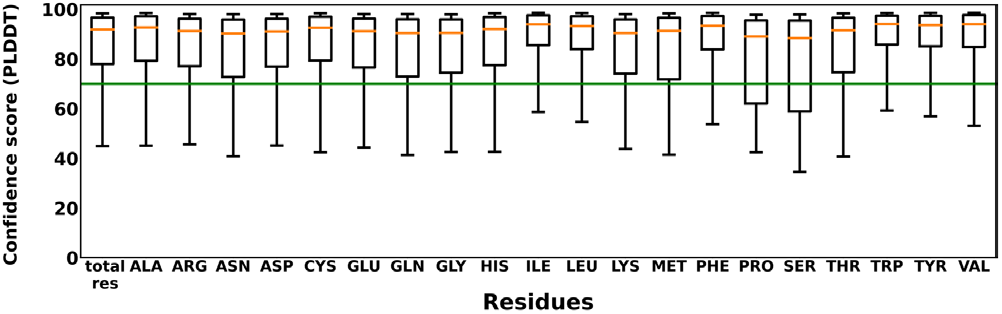
The distribution of PLDDT with the 20 amino acid types in batch 1. Starting from bottom whisker, the boxplots show 10th, 25th, 50th, 75th and 90th percentiles. The horizontal line is at PLDDT = 70 for medium prediction confidence.

The pairwise differences between median prediction confidences provide an additional perspective on the effect of amino acid type on the variation of AlphaFold2’s accuracy. In Figure 6, we present the pairwise differences between PLDDT scores for all amino acids in batch 1. The data for other batches are presented in Figure S14. The maximum pairwise difference of 5.6 occurs between SER and TRP. Another observation is that the highest pairwise differences tend to fall in the PRO and SER columns because AlphaFold2 has the lowest prediction confidence for PRO and SER among all amino acid types.

**Figure 6.**
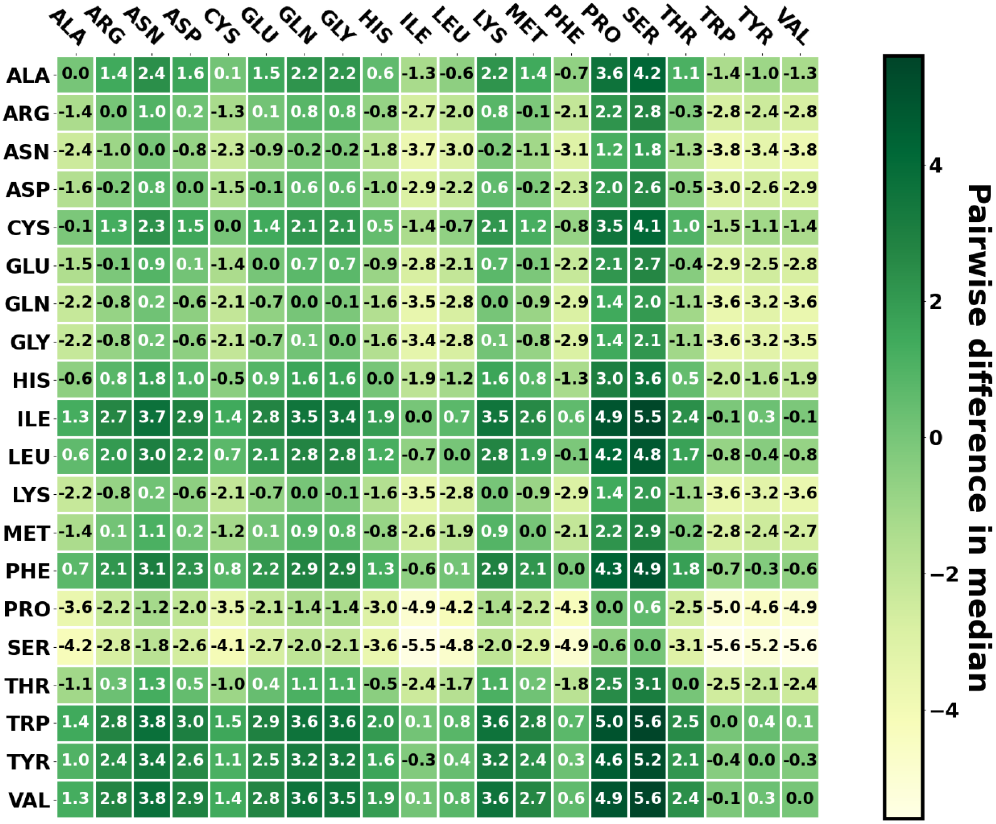
The difference in the median value of PLDDT scores for amino acid pairs in batch 1.

We analyzed the percentage of each amino acid type with PLDDT ≥ 70. In Figure 7, we present these percentages for the 20 amino acids in batch 1. The data for the other batches are presented in Figure S15. Across batch 1, AlphaFold2 meets the medium confidence threshold for at least 70% of each amino acid. We also observe some relative variations when it comes to model fairness for each amino acid. AlphaFold2 has the highest percentage of medium prediction confidence for the following amino acids: TRP (87%), TYR (86%), ILE (86%), LEU (85%), PHE (85%), and VAL (85%). The lowest prediction confidences are observed for SER (70%) and PRO (71%). This agrees with the boxplots for PLDDT distribution in Figure 5.

**Figure 7.**
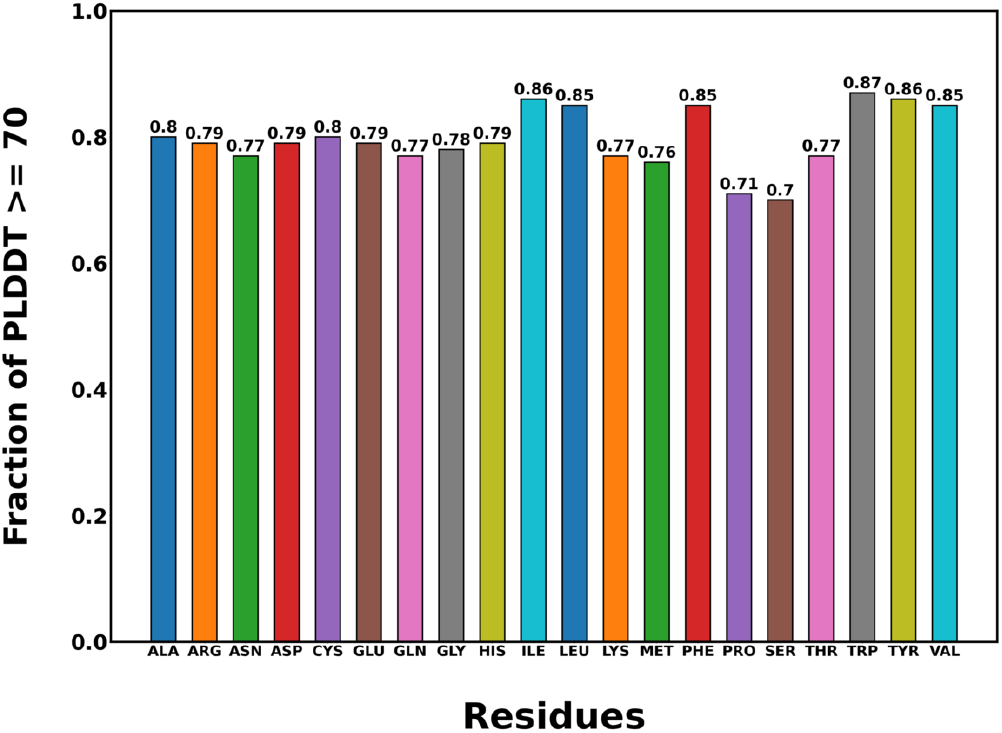
The fraction of each amino acid type with PLDDT ≥ 70 in batch 1.

### 3.3 The distribution of PLDDT with secondary structure

We present the distribution of AlphaFold2’s prediction confidence for secondary structures in batch 1 in Figure 8. Distributions for secondary structures in the other batches can be found in Figure S14. The secondary structures with the highest median prediction confidence are beta-sheet (96.00), beta-bridge (94.56), five-helix (94.38), alpha-helix (93.19) and three-helix (91.38). The worst prediction confidence is observed for coils with the 25th percentile dropping well below 40, which is lower than the criteria for low prediction confidence. This is unsurprising for coils because they are typically disordered structures [34]. Across all the batches, the median prediction confidence for each secondary structure is consistent, with only slight differences in the percentiles.

**Figure 8.**
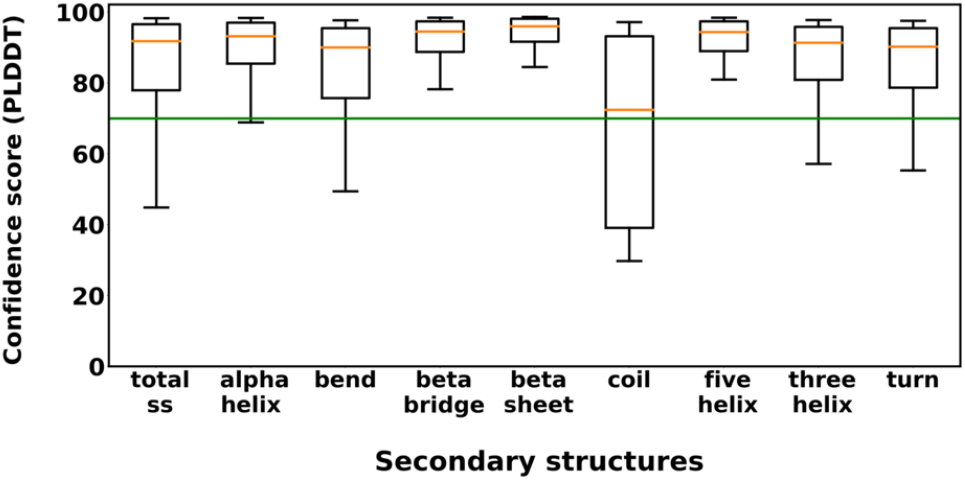
The distribution of PLDDT scores with secondary structure in batch 1.

In Figure 9, we present the pairwise differences between the median PLDDT scores for all secondary structures in batch 1. Pairwise differences for batches 2-5 are presented in Figure S17. The maximum pairwise difference of 23.6 occurs between the beta-sheet and coil. Another observation is that the highest pairwise differences tend to fall in the coil column, which is expected since the lowest prediction confidences are observed for coils. This is consistent with the relatively lower prediction confidences observed for coils in Figure 8.

**Figure 9.**
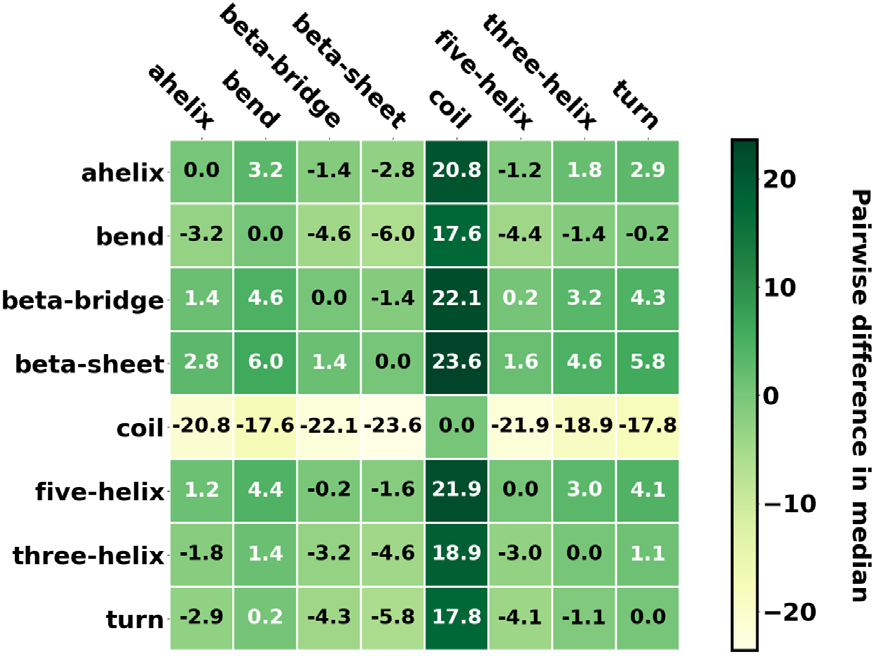
The difference in median value of PLDDT scores between secondary structure pairs in batch 1.

In Figure 10, we present the percentage of each secondary structure with PLDDT ≥ 70 in batch 1. Dats for other batches are presented in Figure S18. In batch 1, AlphaFold2 has PLDDT ≥ 70 for at least 70% of each secondary structure except for coils. Some relative variations are observed when it comes to model fairness with some secondary structures presenting very high percentages of medium prediction confidences while some present relatively lower percentages. AlphaFold2 displays the highest percentage of medium confidence in its beta-sheet, five-helix, and beta-bridge predictions with 96%, 96% and 94% respectively. The worst percentage of medium prediction confidences are seen in coils with AlphaFold2 being confident in only 51% of its predictions. These observations agree with the boxplots in Figure 8.

**Figure 10.**
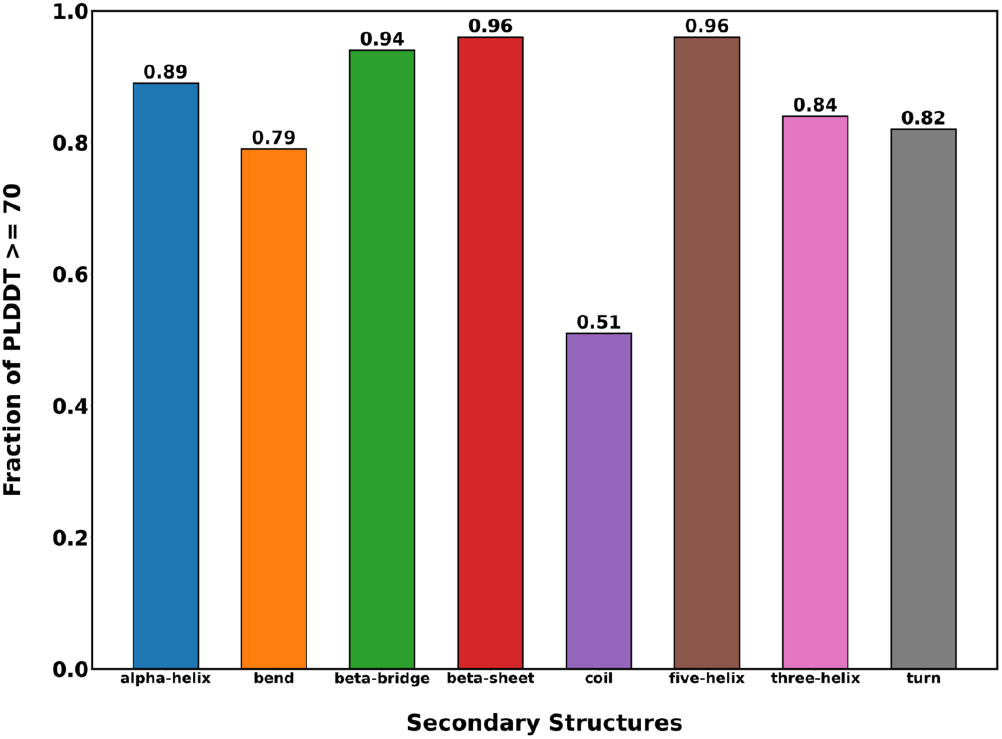
The percentage of each secondary structure with PLDDT ≥ 70 in batch 1.

### 3.4 The influence of protein size on the PLDD distribution with amino acid type and secondary structure

When the PLDDT scores for amino acids are grouped according to protein size N for batch 1, some trends emerge as seen in Figure 11. Data for other batches are presented in Figures S19 – S23. These trends appear consistent for amino acids with high confidence predictions like ILE, as well as for relatively lower confidence predictions like SER. AlphaFold2 performs worse at the extremes which include proteins with N < 100 amino acids, and N > 1000 amino acids as seen in Figure 11 for ILE and SER respectively. It can be observed in Figure 11b that the best AlphaFold2 prediction confidence for SER across all percentiles occurs for proteins with 300 ≤ N < 500 amino acids. As N increases, it is observed that AlphaFold2 becomes progressively less confident in its predictions, with the 25^th^ percentile dropping below PLDDT of 45.

**Figure 11.**
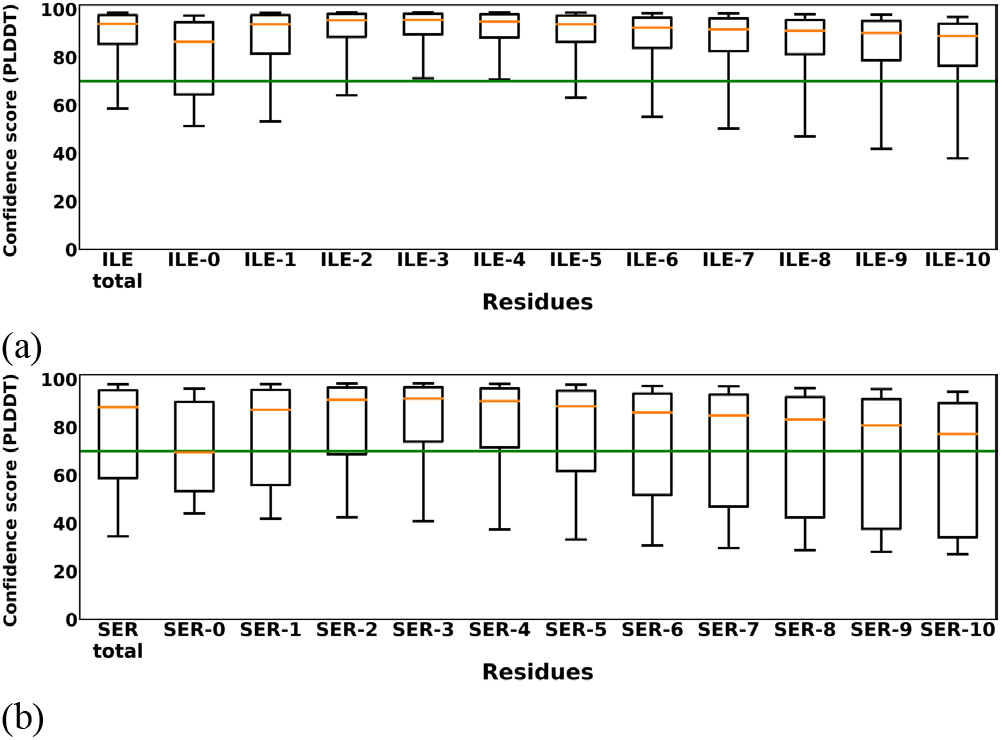
The distribution of PLDDT with the 20 amino acid types as a function of N in batch 1. (a) ILE, and (b) SER. Proteins are grouped in bins of 100 amino acids. SER-0: N < 100 amino acids, SER-1: 100 ≤ N < 200 amino, SER-10: N > 1000 amino acids.

The pairwise differences between median prediction confidences provide an additional perspective to the variation of AlphaFold2’s prediction confidence with protein size. In Figure 12, we present the pairwise differences between the median PLDDT values of amino acids in batch 1. Data from the other batches are presented in Figures S24 – S29. The largest pairwise difference is observed for proteins with N < 100 amino acids, as seen in Figure 12a where the difference is as high as 21.1 between ILE and Methionine for instance. For proteins with 100 ≤ N < 600 amino acids, a progressive decrease to the 4.0 – 6.0 range is observed in the maximum pairwise differences; for instance, the pairwise difference between SER and ILE drops to 5.0 in Figure 12b. As N increases above 600 amino acids, the maximum pairwise difference increases and ranges between 7.0 – 10.0, as can be seen in Figure 12d with the increase of the pairwise difference to 11.6 for SER and ILE. It should also be noted that the PRO and SER columns display the largest pairwise differences because these are the two amino acids with the lowest AlphaFold2 prediction confidences.

**Figure 12.**
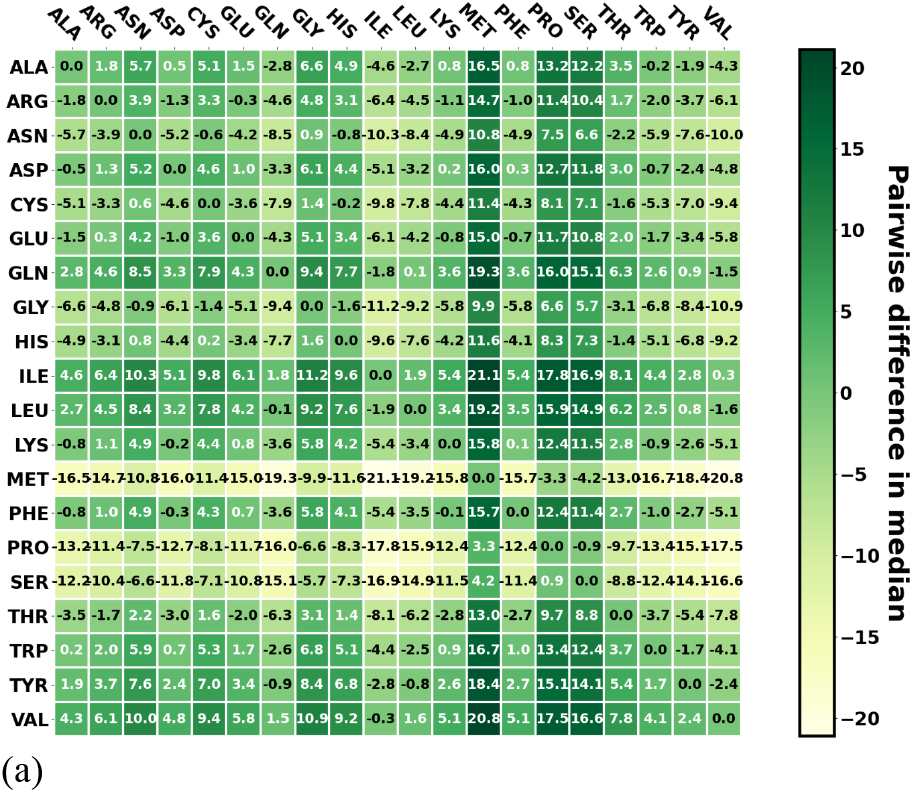
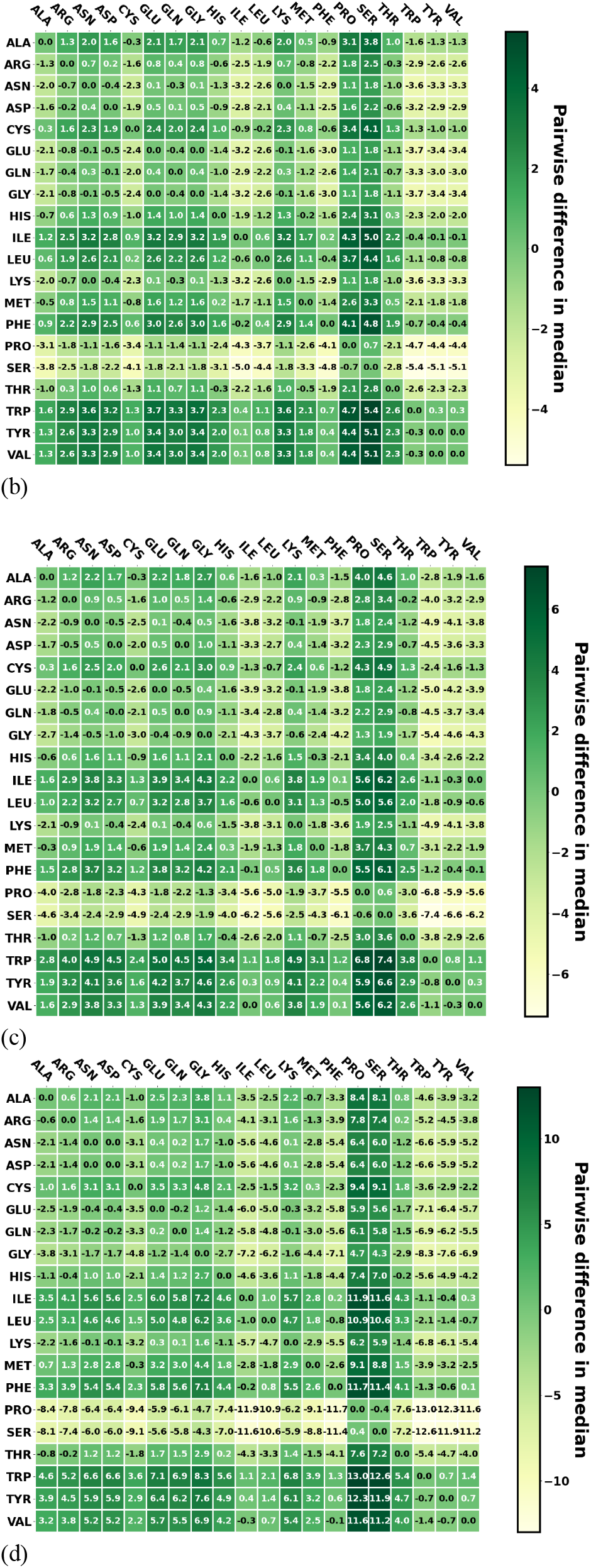
The difference in median value of PLDDT scores for amino acid pairs as a function of protein size N in batch 1. (a) N < 100 amino acids, (b) 500 ≤ N < 600 amino acids, (c) 600 ≤ N < 700 amino acids, (d) N > 1000 amino acids.

Using the accuracy threshold of PLDDT ≥ 70, we present the effects of protein size on the percentage of medium confidence predictions in batch 1 for ILE and SER in Figure 13. The data for batches 2-5 are displayed in Figures S29 – S33. The lowest accuracies are observed proteins with N < 100 amino acids and N > 1000 amino acids. As N increases, the percentage of accurate predictions increases and reaches a maximum for proteins with 300 ≤ N < 400 amino acids. This trend agrees with previous observations.

**Figure 13.**
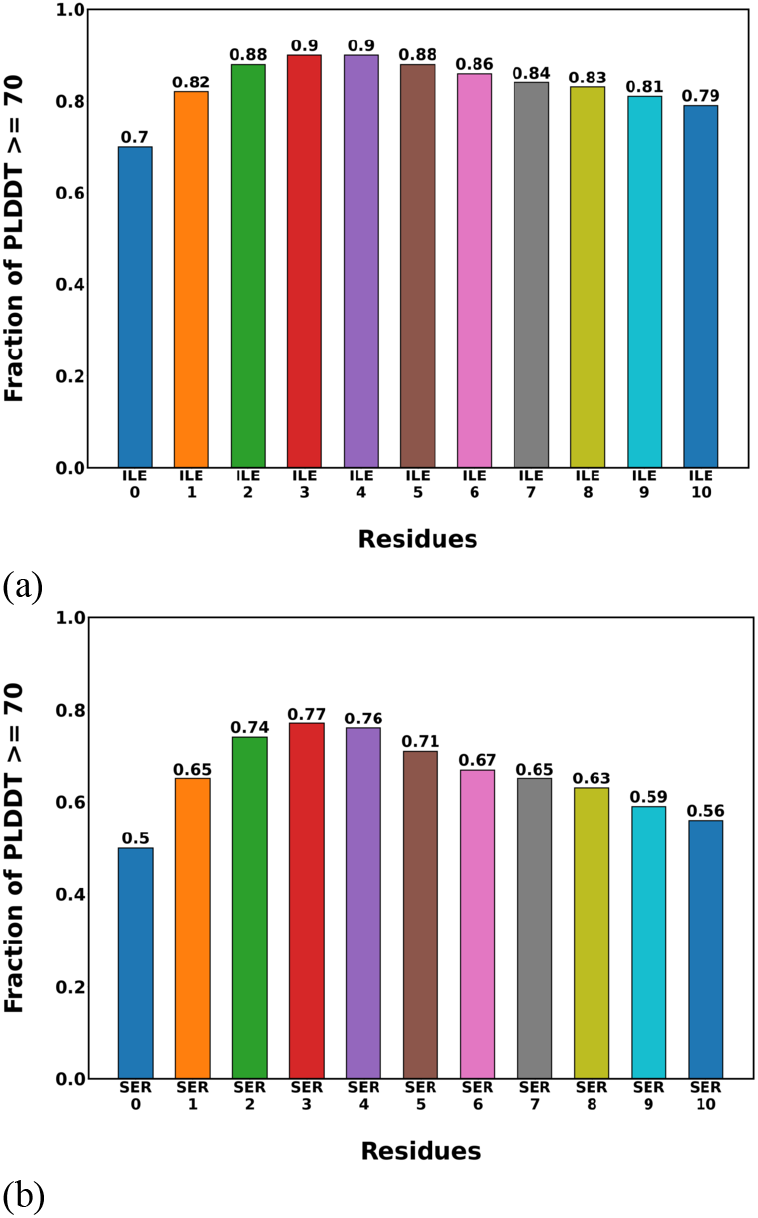
The fraction of each amino acid with PLDDT ≥ 70 as a function of protein size in batch 1. (a) ILE, (b) SER.

When PLDDT scores for secondary structures in batch 1 are grouped according to protein size N, some trends emerge as displayed in Figure 14. Data from batches 2-5 are presented in Figures S34 – S38. With median PLDDT scores above 90, AlphaFold2 displays high confidence in its predictions for alpha-helix, beta-sheet, and beta-bridge. The worst confidence is seen in its coil predictions, with 25^th^ percentile PLDDT scores dropping below 40. Similar to our observation for amino acids, AlphaFold2 performs best at predicting structures of proteins with 300 ≤ N < 400 amino acids as seen in Figure 14b. The worst performances across all percentiles are observed for N < 100 amino acids and N > 1000 amino acids. At N > 400 amino acids, there is a decline in prediction confidence as N increases.

**Figure 14.**
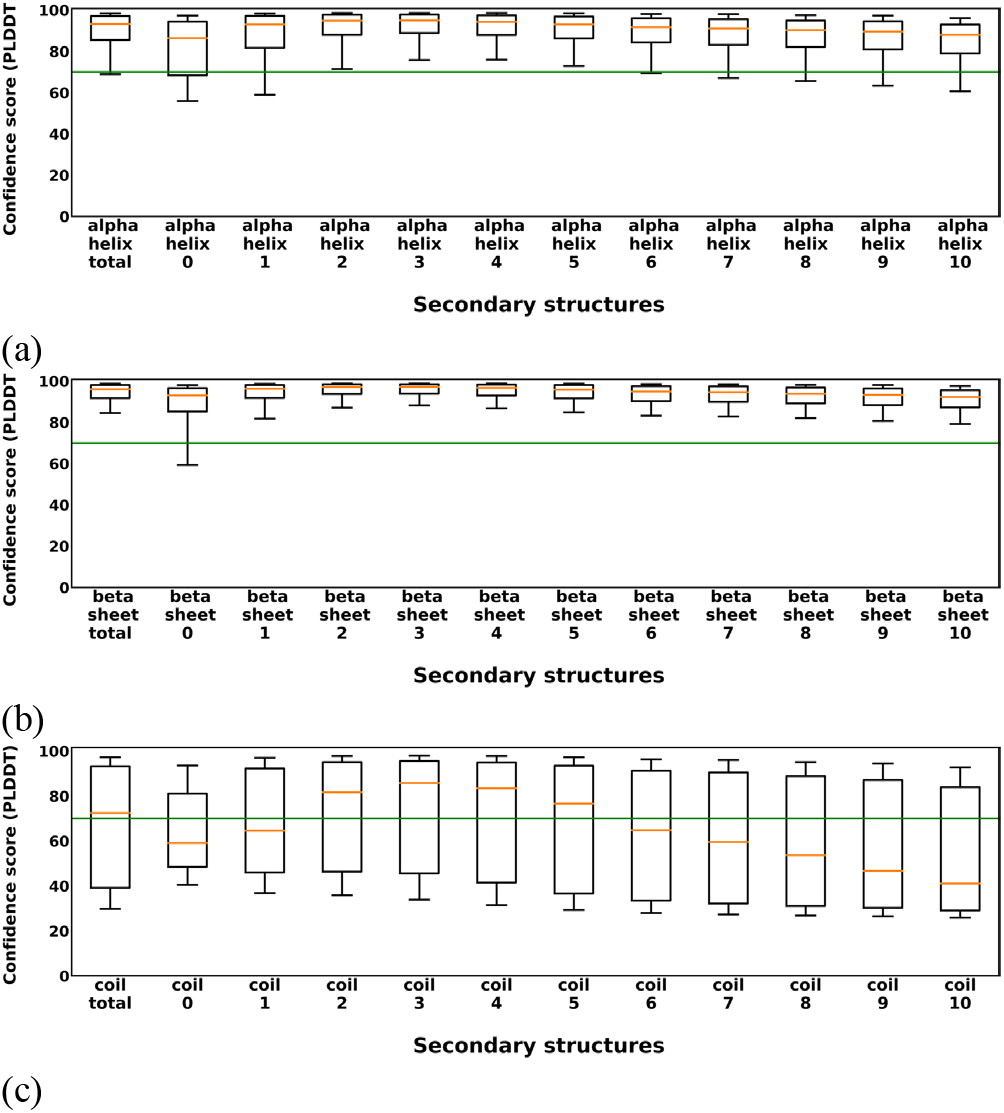
The distribution of PLDDT with the secondary structure as a function of protein size N in batch 1. (a) alpha-helix, (b) beta-sheet, (c) coil. Grouping is done in bins of 100 amino acids. coil-0: N < 100 amino acids, coil-1: 100 ≤ N < 200 amino, coil-10: N > 1000 amino acids.

In Figure 15, we present the pairwise differences in median PLDDT values for secondary structures in batch 1. Batches 2-5 are presented in Figures S39 – S43. The first thing that stands out is how the maximum pairwise differences vary as protein size increases. For proteins with N < 200 amino acids, the maximum pairwise difference is 31.8. For proteins with 200 ≤ N < 600 amino acids, the maximum pairwise difference drops to the 15 – 20 range. At N > 600 amino acids, the maximum pairwise difference jumps back up to 30 and above. The highest pairwise difference is observed for proteins with N > 1000 amino acids. This trend mirrors the observation on AlphaFold2’s performance in Figure 14. The second observation is that the highest pairwise differences tend to fall in the coil column. This is consistent with the relatively lower prediction confidences observed for coils.

**Figure 15.**
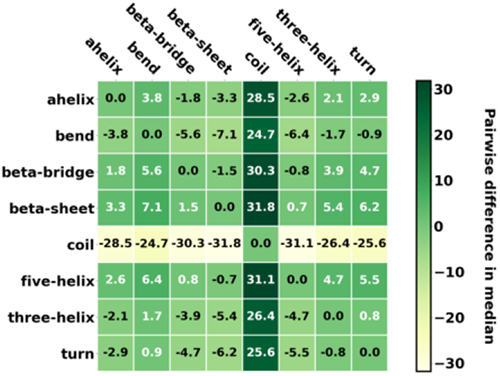
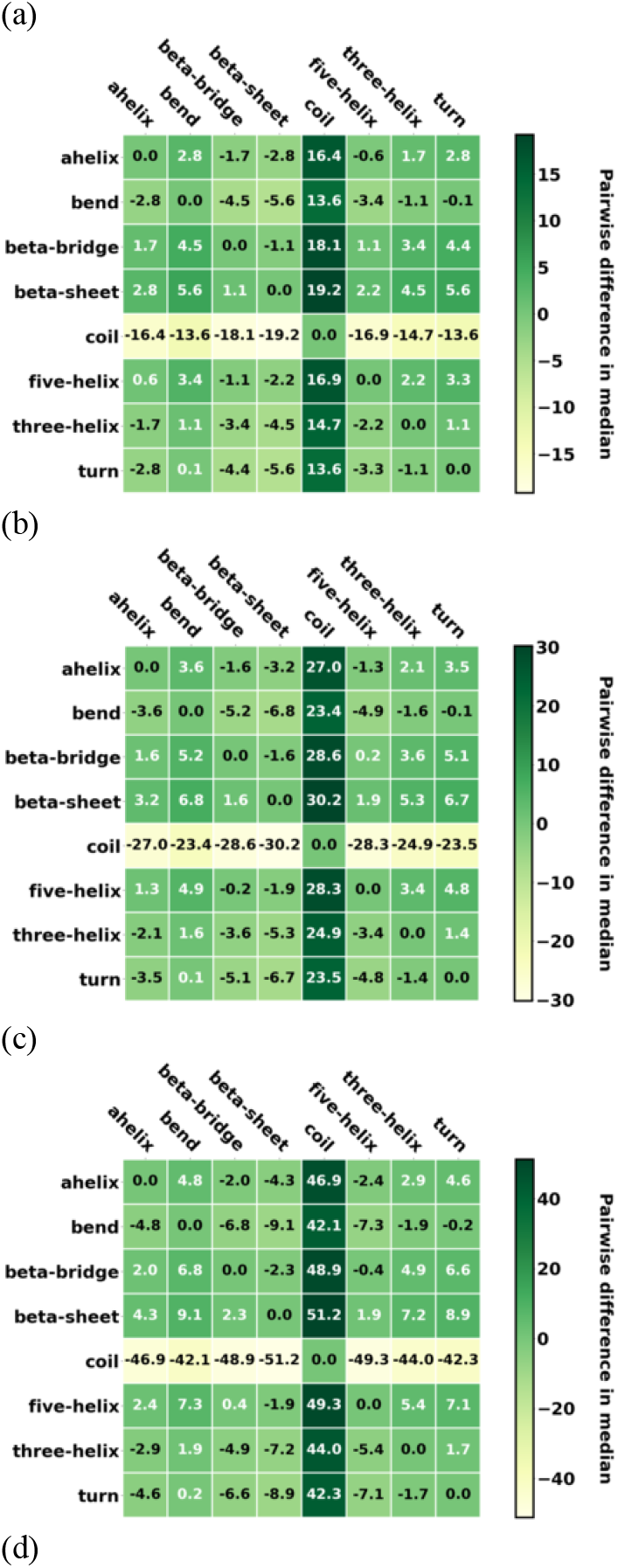
The difference in median value of PLDDT scores between secondary structure pairs as a function of protein size N in batch 1. a) N < 200 amino acids, (b) 500 ≤ N < 600 amino acids, (c) 600 ≤ N < 700 amino acids, (d) N > 1000 amino acids.

Using the threshold of ≥ 70 for PLDDT, we present the effects of protein size on the percentage of accurate predictions in batch 1 for each secondary structure in Figure 16. Data from other batches are presented in Figures S44 – S48. The lowest accuracies are observed for proteins with N < 100 amino acids and N > 1000 amino acids. As N increases, the percentage of accurate predictions increases and reaches a maximum for proteins with 300 ≤ N < 400 amino acids. This trend agrees with previous observations.

**Figure 16.**
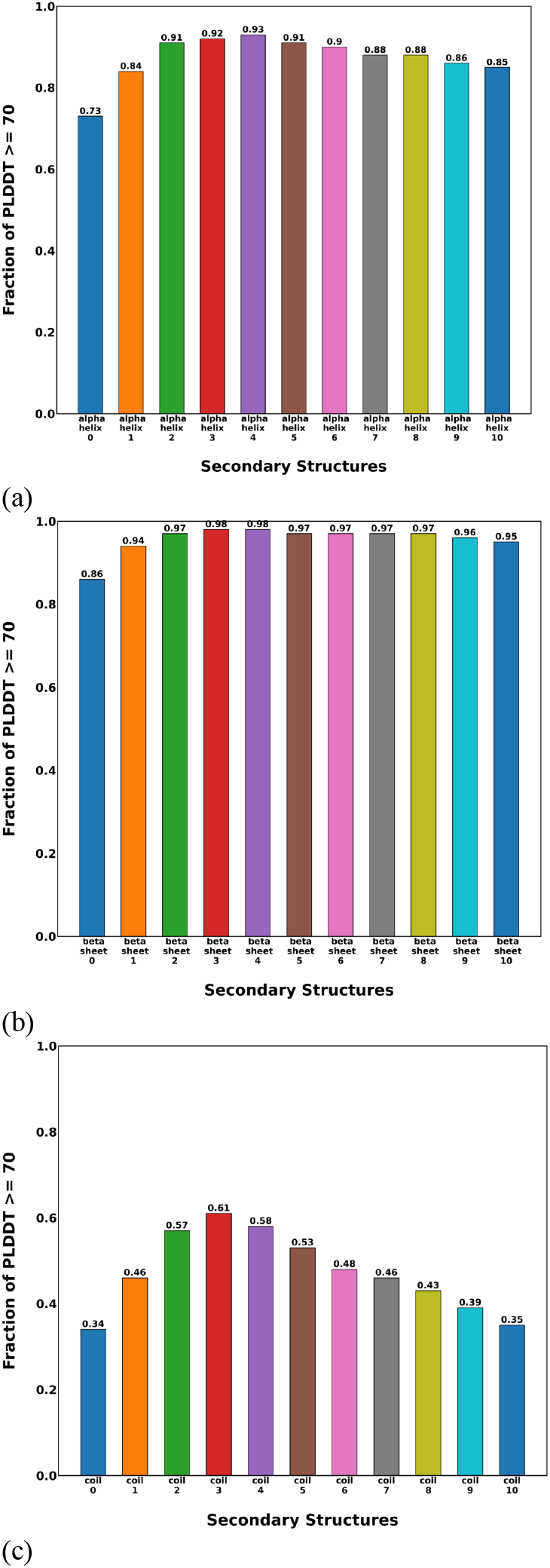
The percentage of each secondary structure with PLDDT ≥ 70 as a function of protein size in batch 1. (a) alpha helix, (b) beta sheet and, (c) coil.

## 4 CONCLUSION

In this work, we carried out a statistical analysis of the prediction confidence of AlphaFold2 across different amino acids, secondary structures, and protein sizes based on 5 million protein structures from the AlphaFold2 repository. From our results, we observe that protein size, amino acid and secondary structure type influence the prediction confidence of AlphaFold2. For amino acids, AlphaFold2 presents the highest prediction confidence for ILE, LEU and TRP; and the lowest prediction confidence for PRO and SER. AlphaFold2 also presents high prediction confidence for secondary structures like beta-sheet, beta-bridge and alpha-helix, and the lowest prediction confidence for coils. Based on protein size, AlphaFold2 displays the highest confidence for predictions on proteins with about 300 – 500 amino acids. The lowest confidence is observed for proteins with less than 100 or greater than 1000 amino acids. In the future, more work needs to be done such as why certain amino acids have low AlphaFold2 PLDDT scores and others do not.

## ACKNOWLEDGMENTS

We would like to thank AI in Medicine (AIM), University of Kentucky (NCATS UL1TR001998, NCI P30 CA177558), and the University of Kentucky Center for Computational Sciences and Information Technology Services Research Computing for their support and use of the Morgan Compute Cluster and associated research computing resources.

